# Predicted structural mimicry of spike receptor-binding motifs from highly pathogenic human coronaviruses

**DOI:** 10.1101/2021.04.23.441187

**Authors:** Christopher A Beaudoin, Arian R Jamasb, Ali F Alsulami, Liviu Copoiu, Andries J van Tonder, Sharif Hala, Bridget P Bannerman, Sherine E Thomas, Sundeep Chaitanya Vedithi, Pedro H M Torres, Tom L Blundell

**Author notes:** corresponding author and lead contact Joint corresponding authors email addresses: Tom Blundell, Chris Beaudoin.

## Abstract

Viruses often encode proteins that mimic host proteins in order to facilitate infection. Little work has been done to understand the potential mimicry of the SARS-CoV-2, SARS-CoV, and MERS-CoV spike proteins, particularly the receptor-binding motifs, which could be important in determining tropism of the virus. Here, we use structural bioinformatics software to characterize potential mimicry of the three coronavirus spike protein receptor-binding motifs. We utilize sequence-independent alignment tools to compare structurally known or predicted three-dimensional protein models with the receptor-binding motifs and verify potential mimicry with protein docking simulations. Both human and non-human proteins were found to be similar to all three receptor-binding motifs. Similarity to human proteins may reveal which pathways the spike protein is co-opting, while analogous non-human proteins may indicate shared host interaction partners and overlapping antibody cross-reactivity. These findings can help guide experimental efforts to further understand potential interactions between human and coronavirus proteins.

**Highlights:**

- Potential coronavirus spike protein mimicry revealed by structural comparison
- Human and non-human protein potential interactions with virus identified
- Predicted structural mimicry corroborated by protein-protein docking
- Epitope-based alignments may help guide vaccine efforts

**Graphical abstract:** 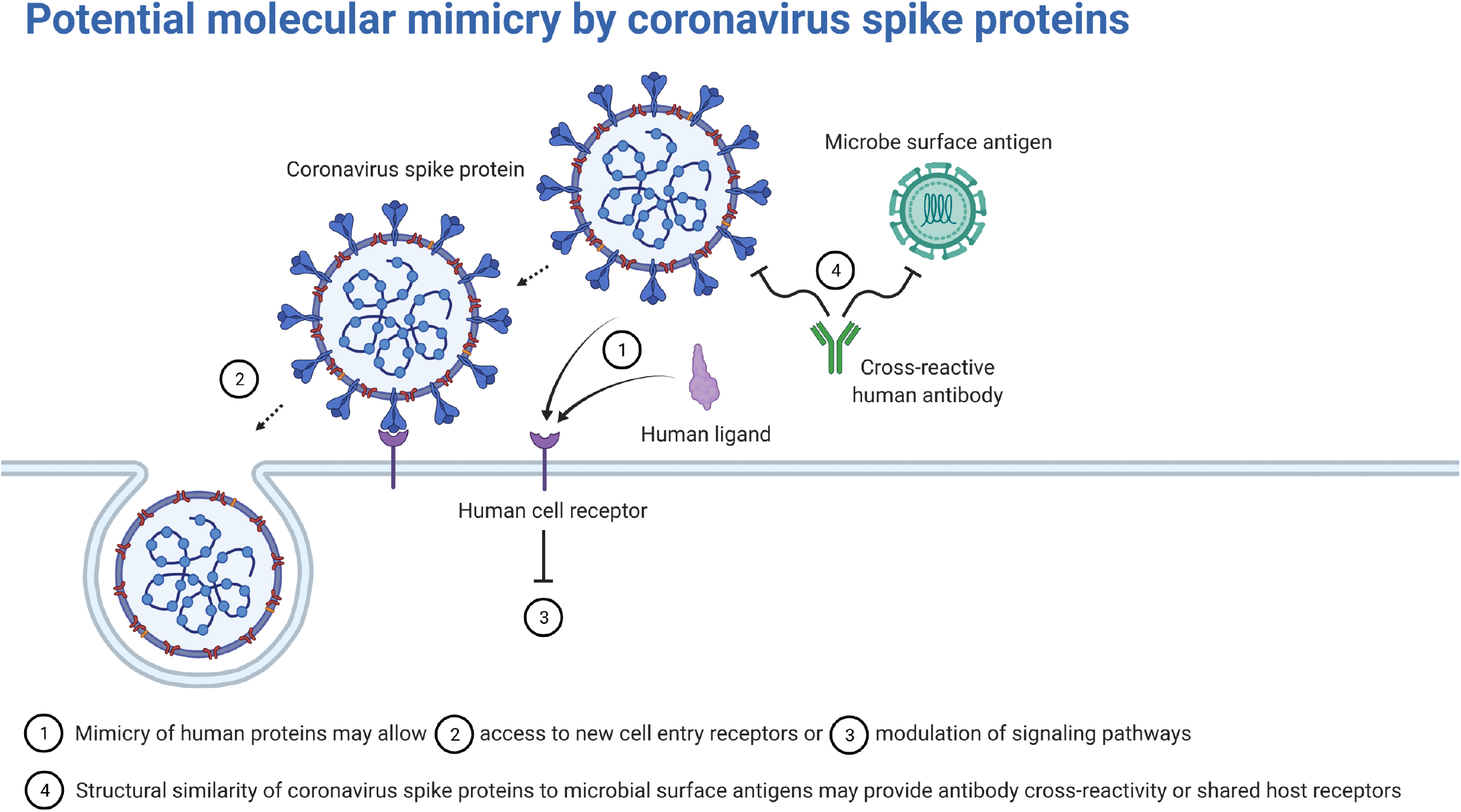

## Introduction

Viruses have long been known to utilize molecular mimicry of host proteins to interrupt and exploit host biochemical pathways during infection (Alcami, 2003; Lasso et al., 2021). Alongside the need to employ host machinery for the viral replication cycle, the evolution of viral protein motifs that resemble host proteins can result in new virulence mechanisms, such as inducing inflammation and evading the immune system (Elde and Malik, 2009). Coronaviruses, in particular, have been suspected to have acquired human protein mimics throughout the long record of human coronavirus infections (Chew et al., 2003; Walls et al., 2019). As further evidence, the highly pathogenic human coronaviruses, Severe Acute Respiratory Syndrome Coronavirus 2 (SARS-CoV-2), SARS-CoV, and the Middle Eastern Respiratory Syndrome Coronavirus (MERS-CoV), have been shown to encode numerous short linear motifs across their genomes that are homologous to human proteins (Martínez et al., 2021). Although coronavirus infections are typically localized to the lungs, resulting in respiratory infections, viral material has also been found in other organs, such as the kidney, brain, and heart, resulting in more life-threatening infections (Renu et al., 2020). Furthermore, SARS-CoV-2 (the causative agent of the COVID-19 pandemic) infection has presented symptoms not previously seen in other coronavirus infections, such as conjunctival discharge from the eyes (Cheong, 2020; Zheng et al., 2021). Investigations into coronavirus host mimicry may shed light on viral tropism and infection severity (Angileri et al., 2020a).

The structure of the receptor-binding motif (RBM) on the spike glycoprotein is particularly important for determining the tropism of the virus (Reguera et al., 2014). Host receptors that contain motif(s) that complement the electrochemical and spatial configurations of the viral RBM will interact and, thus, initiate viral entry (Li, 2015; Tortorici and Veesler, 2019). Angiotensin converting enzyme II (ACE2) has been established as the primary cell entry receptor for SARS-CoV-2 and SARS-CoV and dipeptidyl peptidase IV (DPP4) as the primary cell entry receptor for MERS-CoV. However, several reports, some preliminary, have proposed additional coronavirus cell entry receptors, such as transferrin receptor protein 1, kidney injury molecule-1, kremen protein 1, and αv integrins for SARS-CoV-2 (Gu et al., 2020; Ichimura et al., 2020; Li et al., 2003; Sigrist et al., 2020; Tang et al., 2020; Wang et al., 2013; Yang et al., 2020). Additionally, coronavirus spike proteins have been proposed to interact with host factors to facilitate infection aside from their role in cell entry (Patra et al., 2020). For instance, two studies found that the SARS-CoV-2 spike protein alone can interact with the blood brain barrier (Buzhdygan et al., 2020; Rhea et al., 2021). The importance in receptor-binding and low glycosylation surrounding the coronavirus RBM residues make it an attractive target for inhibition by small-molecule drugs, therapeutic peptides, and neutralizing antibodies (Hussain et al., 2020; Ling et al., 2020; Pandey et al., 2020).

To date, there has been limited investigation into the structural similarity of highly pathogenic coronavirus RBMs (Kanduc and Shoenfeld, 2020). Identifying structurally analogous human proteins may give insight into endogenous biochemical pathways that the virus is hijacking to facilitate infection or may help explain autoimmune disorders triggered by coronavirus infections (Angileri et al., 2020b; Drayman et al., 2013). Detecting similar microbial proteins may reveal shared host receptors or antibody cross-immunity (Huang et al., 2020). Short linear motifs on coronavirus spike RBMs have been shown to share high amino acid sequence identity with human proteins, which may indicate host mimicry (An and Park, 2020; Grifoni et al., 2020; Lin et al., 2020; Lucchese and Flöel, 2020). However, protein structure and fold similarity have been shown as more informative than amino acid sequence similarity in predicting molecular mimicry (Krishna and Grishin, 2004; Westall, 2006). Drayman *et al*. performed a structural similarity search using bacterial and viral motifs and experimentally validated the simian vacuolating virus 40 major capsid protein mimicry of Gas6 binding with TAM – Tyro3, Axl, and Mer – receptors, demonstrating that structural paralogs with low amino acid identity may still act as molecular mimics. Thus, to add to the understanding of host mimicry of highly pathogenic coronavirus RBMs, we used structural bioinformatics tools to model and map the extent to which the three-dimensional structures of the SARS-CoV-2, SARS-CoV, and MERS-CoV spike RBMs are potentially mimicking the interactions of experimentally-determined protein structures. We used structural alignment tools with distinct methodologies to perform a structural similarity screen between the RBMs and all known protein structures and, subsequently, tested potential RBM interactions with protein-protein docking simulations. Several cell signaling proteins, innate immune factors, snake and spider toxins, and microbial antigens are found to share structural features with the three RBMs. This information may help guide experimental efforts to elucidate spike RBM interactions, including that of vaccine design and cell entry receptor discovery.

## Results and Discussion

### Receptor-binding motif structural similarities and characteristics

Several models of the spike protein for each of the highly pathogenic coronaviruses have been experimentally determined; however, many of them are missing residues due to the difficulty in resolving the structure of flexible protein motifs (Nwanochie and Uversky, 2019). To overcome this issue and obtain a representative three-dimensional model of each spike receptor-binding motif (RBM), we used ProtCHOIR, a recently developed pipeline to automate the modelling of homo-oligomers, to model each trimeric spike protein and, subsequently, manually selected the RBM residues for each coronavirus (Figure 1). All generated models were structurally aligned to experimental models using TM-align to determine modelling precision. On a scale from 0 to 1, a TM-score of over 0.5 between two proteins implies that they have the same fold, while below 0.2 suggests a random alignment. Each RBM alignment with the corresponding experimental structure reported a TM-score over 0.95, reflecting high-quality modelling. Although receptor-binding of coronavirus spike proteins has been shown to be an elaborate process that involves interactions with glycans and multiple protein domains, we selected the most interactive region of the spike RBD with primary receptors (i.e. ACE2 for SARS-CoV and SARS-CoV-2; DPP4 for MERS-CoV) from experimental models as the receptor-binding motif (RBM) (Wang and Xiang, 2020).

**Figure 1.**
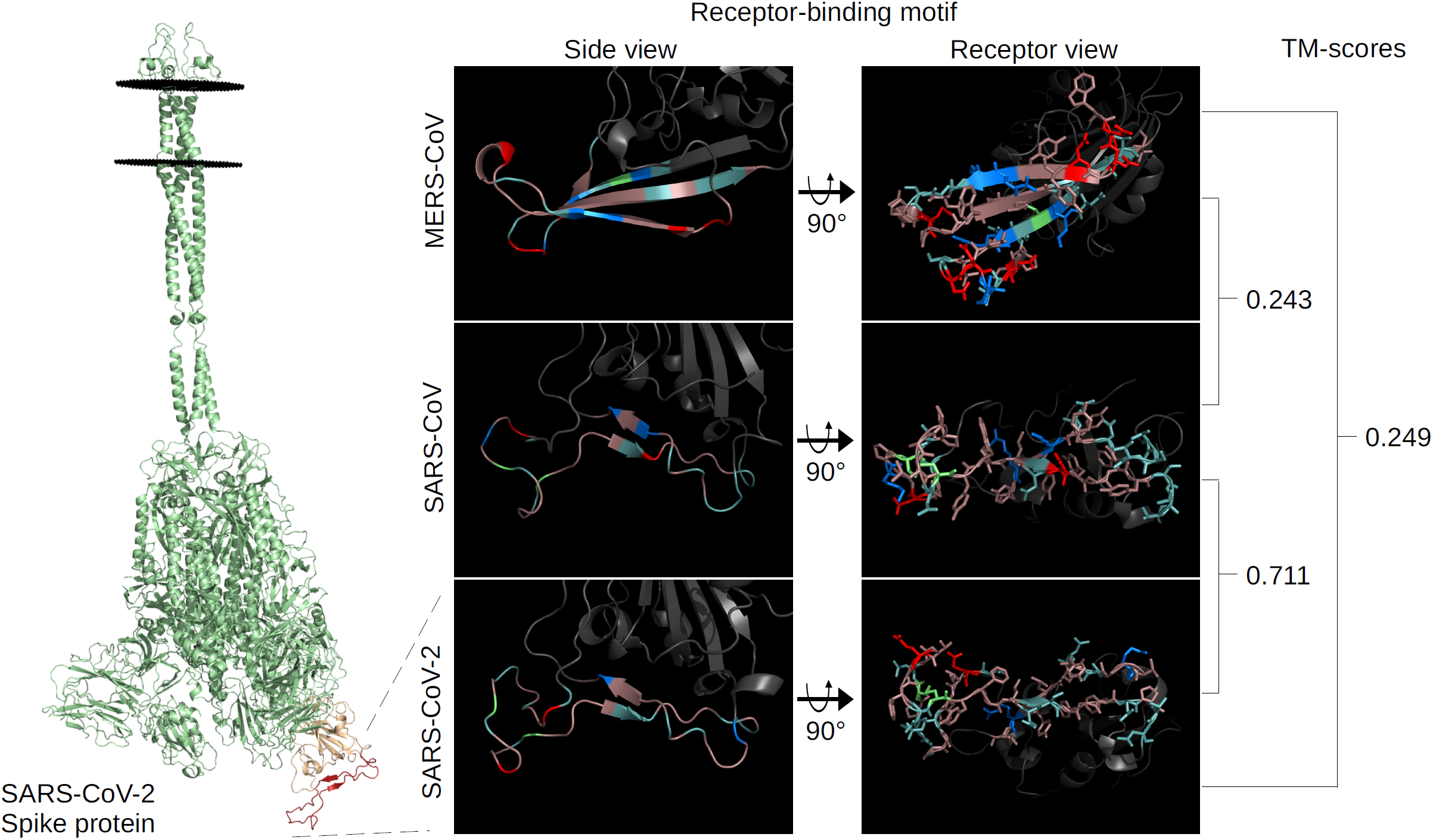
Spike receptor-binding motif comparison. The full-length SARS-CoV-2 spike protein (green), left, modelled using ProtCHOIR is shown with the receptor-binding domain (yellow) and receptor-binding motif (red) marked. The RBMs from the side view are shown, middle, with the amino acids labelled by color: red for acidic (D,E), blue for basic (H,R,K), light teal for polar non-charged (S,N,T,Q), dirty violet for hydrophobic (A,V,I,L,M,F,W,P,G,Y), and lime green for cysteine residues. RBMs from the host cell receptor side are shown, right, with amino acid stick configurations.

The structural similarity of the RBMs to one another was quantitatively assessed using TM-align before assessing their similarity to other known proteins. The SARS-CoV-2 and SARS-CoV RBMs were very similar with a TM-score of 0.71, while the TM-scores of MERS-CoV with the other two were both less than 0.25 (Figure 1). This level of divergence is also reflected at the amino acid sequence level for the RBD of SARS-CoV-2 and SARS-CoV at 64.6% sequence identity and MERS-CoV with SARS-CoV-2 and SARS-CoV at 19% and 21.6%, respectively.

As seen in Figure 1, the SARS-CoV-2 RBM is comprised almost exclusively of hydrophobic and polar non-charged amino acids, with the exception of one acidic glutamate and one basic lysine. SARS-CoV is similar to SARS-CoV-2 in that it is composed mostly of hydrophobic and polar non-charged residues with some exceptions as single amino acid differences, such as an acidic aspartate in the middle of the SARS-CoV RBM. The MERS-CoV RBM consists of more acidic and basic amino acids and contains fewer polar non-charged residues. Of note, the SARS-CoV-2 and SARS-CoV RBMs have 7 and 8 aromatic residues, respectively, exposed on the receptor-binding surface of the RBM. The recent discovery of the N501Y and E484K mutants add a potentially functional aromatic and basic residue, respectively, in the SARS-CoV-2 RBM – both of which have been proposed to increase binding to ACE2 (Nelson et al., 2021). Modelling of the mutants yielded very small structural changes in the SARS-CoV-2 RBM – TM-scores of the mutant RBMs aligned to the reference structure were above 0.9.

In terms of global architecture, the SARS-CoV-2 and SARS-CoV RBMs contain two anti-parallel beta-strands connecting three loops, although the SARS-CoV-2 RBM has two short beta-strands leading to a cystine disulfide loop (Figure 1). Both SARS-CoV and SARS-CoV-2 contain a similar cystine disulfide bond helping shape one end of the respective RBMs. The MERS-CoV RBM consists of three beta-strands connecting four loops. Because loop flexibility may affect overall structure, we submitted each RBD to the CABS-flex 2.0 web server and found that the cystine disulfide loop of both the SARS-CoV-2 and SARS-CoV RBMs displayed high flexibility (> 9 RMSF) -otherwise, the RBM residues on all three RBMs were predicted to exhibit low RMSF (< 6.5) (Supplementary Figure 1). The flexibility predictions from CABS-Flex 2.0 were supported by separate studies on coronavirus RBMs (Saputri et al., 2020; Spinello et al., 2020). The high flexibility of the cystine loops in the SARS-related RBMs motivated the use of two additional models provided by CABS-Flex 2.0 for the structural similarity screen. The added models reported surprisingly low TM-scores compared to the references (0.42 and 0.65 for SARS-CoV-2 and 0.45 and 0.41 for SARS-CoV), revealing the high flexibility in these loops (Supplementary Figure 1). Overall, SARS-CoV-2 and SARS-CoV were found to share higher structural homology with one another than in comparison with MERS-CoV.

### Structural similarity screen

After RBM model generation, we performed a structural similarity screen for each RBM. Four sequence-independent 3D-structure alignment tools with different methodologies were used to quantify the structural similarity between the RBMs and known 3D protein structures in order to better understand shared structural features between the RBMs and potential mimics. Notably in this study, although spike may engage in interactions within human cells, we focused on protein structures that would be found in the extracellular matrix (excluding antibodies, due to their structural diversity) to gain more insight into potential cell entry receptors, immunopathies, and shared antigenicity with other microorganisms (Versteeg et al., 2007).

The PDBeFold, RUPEE, and HMI-PRED web servers were used, and TM-align was locally-installed and run pairwise against the downloaded PDB database clustered at 100% sequence identity. The TM-score distributions between SARS-CoV-2 and SARS-CoV were quite similar, while MERS-CoV was more similar to a greater number of proteins (Figure 2A). The MERS-CoV RBM returned 3,954 structures with a TM-score of over 0.5 (∼ top 1% of TM-scores) out of 245,055 total RBM-chain alignments and an average TM-score of 0.33. The SARS-CoV-2 and SARS-CoV RBMs had lower average TM-scores, 0.298 and 0.297 respectively, and the top 1% corresponded roughly to the 0.4 TM-score line. Thus, structures with a TM-score of > 0.4 were selected for further analysis for the SARS-related viruses: 4,025 for SARS-CoV-2 and 3,561 for SARS-CoV. PDBeFold returned 621-806 and 1,163 structures for the SARS-related and MERS-CoV RBM models, respectively. The top 1,000 hits from each RUPEE run were recorded. HMI-PRED outputs ranged from 20-50 mimicked PDB templates per RBM. All alignments of interest were manually inspected to validate the potential for structural mimicry. Returned aligned proteins from each tool were linked to their corresponding PDB and UniProt codes. Shared UniProt codes between two or more tools were regarded as high-confidence hits. Biologically relevant structural alignments specific to each tool were also inspected and considered. Structural alignments that would not make sense biologically, such as when the RBM is facing the inside of the protein, were discarded, while alignments that were logical but found outside of protein-protein interfaces were included on a case-by-case basis. Returned structures not shown to be found in the extracellular matrix were removed. All tools returned their respective spike structures, confirming their validity.

**Figure 2.**
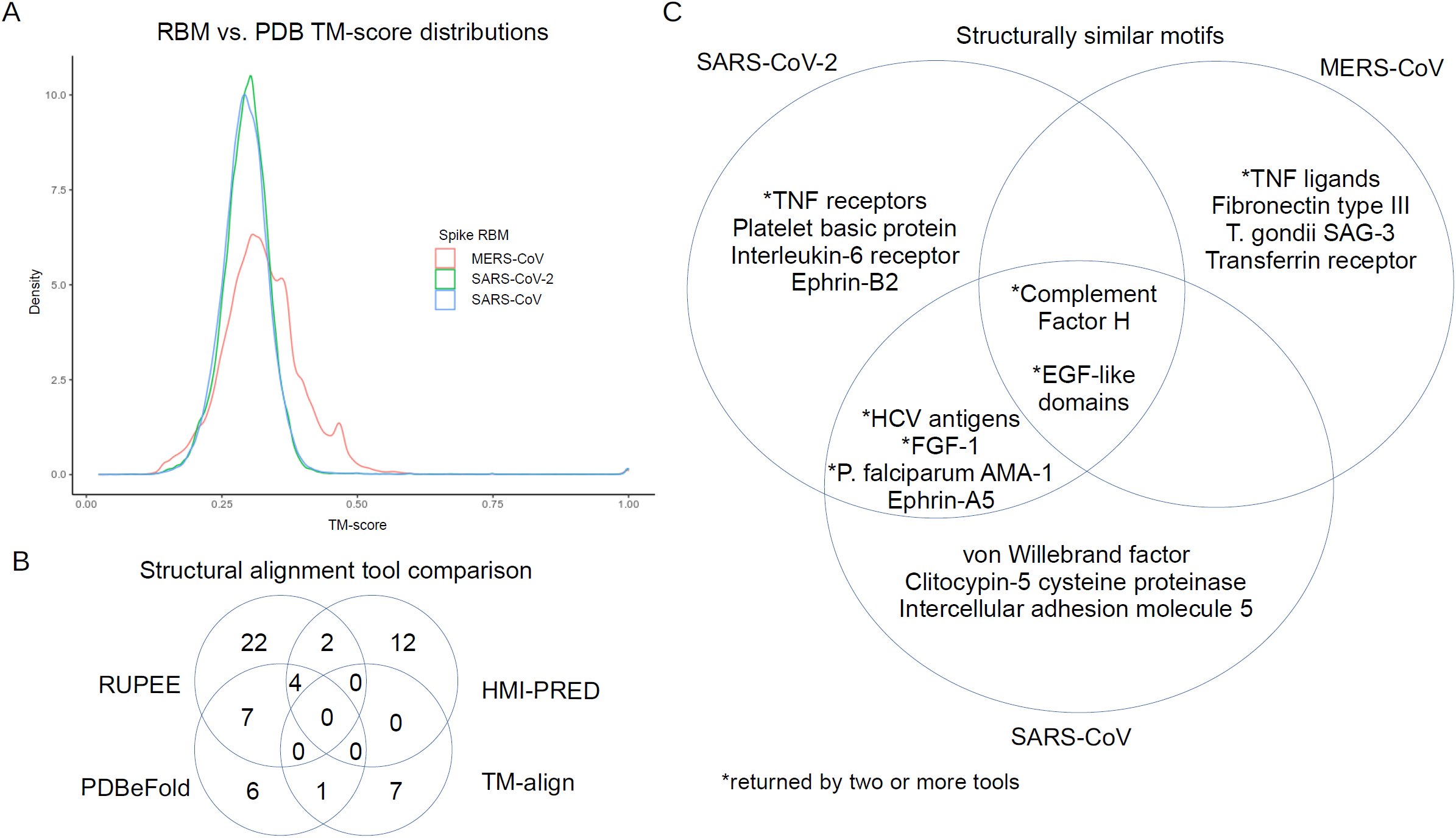
Summary of structural similarity screen. The TM-scores generated from the in-house TM-align screen are displayed as a density plot for each RBM (A). Number of shared proteins from the tools used in the structural similarity screen compared and contrasted (B). Structurally similar motifs, common between coronavirus receptor-binding motifs, compared and contrasted (C).

A total of 62 UniProt codes, excluding 28 toxins, were considered as biologically relevant, which were comprised of 35, 19, 19, and 8 selected alignments from RUPEE, HMI-PRED, PDBeFold, and TM-align, respectively. When comparing tools (Figure 2B), RUPEE and PDBeFold web servers shared 7 UniProt codes for at least one RBM, while TM-align shared 1 with PDBeFold and 0 with RUPEE. HMI-PRED shared 2 structures with RUPEE, 1 with PDBeFold, and 0 with TM-align. Little overlap was shown between most of the tools, which is consistent with structural similarity-based studies on HIV and human proteins (Doolittle and Gomez, 2010). The combined returned UniProt codes, excluding toxins, from all four tools totalled 39, 23, and 29 for the SARS-CoV-2, SARS-CoV, and MERS-CoV RBMs, respectively. The top alignments consisted of cytokines, chemokines, and growth factors and their receptors; structures containing EGF-like domains; complement activation proteins; cystine disulfide-rich toxins derived from snakes and spiders; and antigenic microbial proteins. A Venn diagram showing some shared hits between the three RBMs can be seen in Figure 2C, and a full listing of the hits, alignment values, and tools can be found in Tables 1 and 2. The SARS-CoV and SARS-CoV-2 RBMs shared more structural domains, while MERS-CoV returned more unique hits compared to the other two. Altogether these results indicate that proteins from completely different protein families may interact with coronavirus spike RBMs.

**Table 1.**
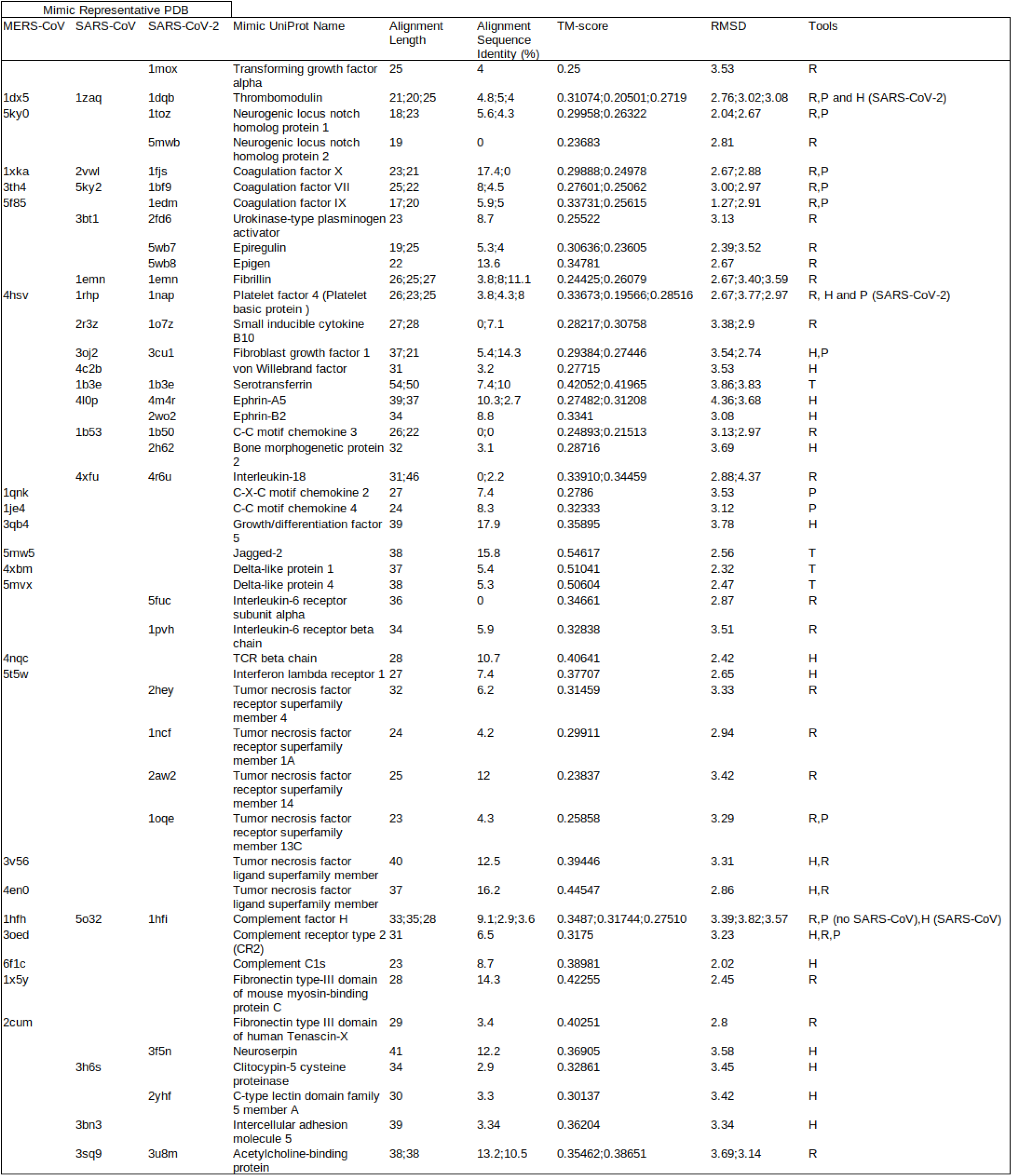
Structural alignment values and data for endogenous hits

**Table 2.**
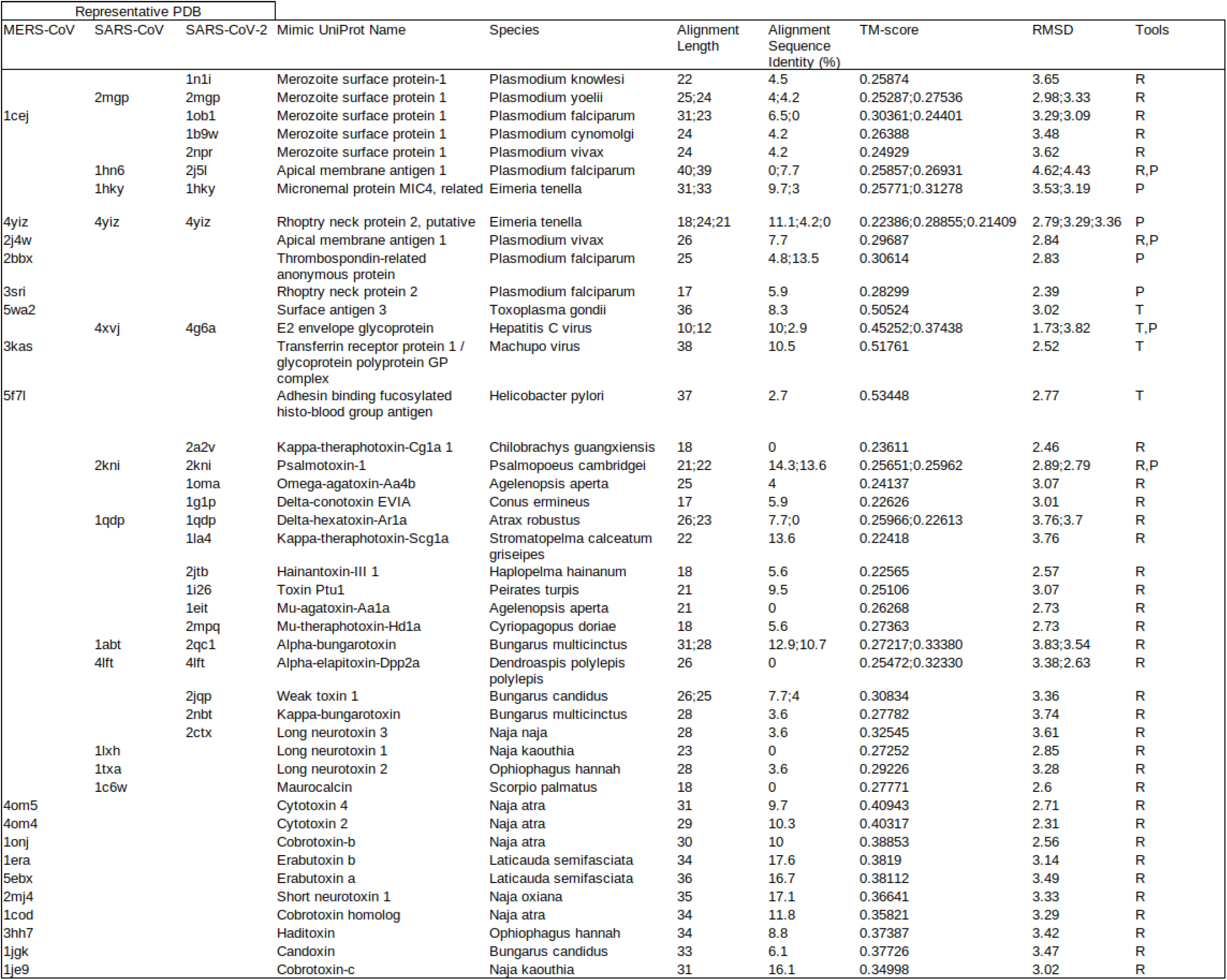
Structural alignment values and data for exogenous hits

### Analysis of predicted structural mimicry

Further examination of the structural alignments and their relevance to biological activity was performed to elucidate potential mechanisms of molecular mimicry by the SARS-CoV-2, SARS-CoV, and MERS-CoV spike RBMs. The UniProt and STRING databases were used to link the predicted mimics with potential interaction partners, and the PDB provided template structures to determine whether the alignments were found in ligand-binding regions. Selected high-confidence potential interactions were further evaluated using protein-protein docking with ClusPro PIPER in order to better understand electrochemical, in addition to structural, complementarity considering the low amino acid sequence identity. The docked models were then analyzed with the FoldX AnalyseComplex program to determine the complex interaction energy. Docking of the natural ligand to the receptor was performed to obtain a control interaction energy. The energy of the original PDB protein complex was also predicted as an experimental control. The exploration of these interactions with structural alignment visualization and protein-protein docking may help explain their potential roles in infection.

The potential mimics were split into two categories: endogenous vs. exogenous, or human vs. non-human, to more effectively describe the results in the context of infection. Mimicry of endogenous proteins may reveal which human pathways, specifically, the viral RBM is hijacking; structurally similar exogenous proteins may exhibit shared interference of human interaction pathways or antigenicity with the coronavirus RBMs. Endogenous hits, both discovered by single and multiple structural alignment tools, are summarized in Table 1 and exogenous hits in Table 2.

#### Endogenous

Several proteins containing EGF-like domains were found to be similar to all three RBMs. EGF-like domains are evolutionarily conserved domains that share homology to the epidermal growth factor and have been shown to function primarily in tissue organization and repair (Engel, 1989; Tombling et al., 2020). Both the cystine disulfide loop and the central beta-strand sub-motif structures in the SARS-CoV-2 and SARS-CoV RBMs and the MERS-CoV beta-strands were found to mimic EGF-like domains.

The EGF-like domain of the urokinase-type plasminogen activator (uPa) in complex with its receptor, urokinase plasminogen activator receptor (uPAR), (PDB: 2fd6) was found to be similar to both SARS-CoV-2 and SARS-CoV RBMs using RUPEE. Interestingly, the uPa/uPAR system has been implicated in SARS-CoV-2 pathogenesis with uPAR as an early predictor of severe respiratory failure (D’Alonzo et al., 2020; Rovina et al., 2020). Although the RBMs protrude into the receptor in the structural alignments, the alignments suggest that the RBMs might bind to uPAR (Supplementary Figure 2A).

The neurogenic locus notch homolog protein 1 (NOTCH1) EGF-like domain was returned for the SARS-CoV-2 RBM central beta-strands and MERS-CoV RBM by RUPEE and PDBeFold. NOTCH1 is involved in developmental, innate immunity, and inflammation signaling pathways, and natural ligands of the NOTCH1 EGF-like domains include jagged-1, jagged-2, delta-like 1 (DLL1), DLL3, and DLL4 (Shang et al., 2016). Alignment of the SARS-CoV-2 and MERS-CoV RBMs with the EGF-like domain of NOTCH1 bound to DLL4 (PDB: 4xl1) shows potential for molecular mimicry, i.e. the coronavirus RBMs may bind to DLL4 (Supplementary Figure 2B) (Luca et al., 2015). The SARS-CoV-2 RBM was also found similar to NOTCH2 by RUPEE, but no PDB complex models were available for further inspection. No direct interactions with the NOTCH1 pathway have been revealed, but its inhibition has been proposed to help fight SARS-CoV-2 infection (Rizzo et al., 2020).

All three RBMs were found to potentially mimic the EGF-like domain of coagulation factor VIIa. Further inspection of the alignment in complex with tissue factor (PDB: 1dan) showed potential for mimicry (Supplementary Figure 2C) (Banner et al., 1996). Interestingly, tissue factor expression has been shown to be up-regulated in severe SARS-CoV-2 infections, although there are several plausible theories (Bautista-Vargas et al., 2020; Eslamifar et al., 2020). The cystine disulfide loops of SARS-CoV and SARS-CoV-2 were found to resemble the EGF-like domains of coagulation factors X and IX and fibrillin, which are known to bind calcium (Handford et al., 1995; Stenflo et al., 2000). However, there is no evidence for calcium binding to the RBMs.

All three RBMs were found to mimic the EGF-like domain of thrombomodulin, specifically in the region that binds thrombin (PDB: 1dx5), by RUPEE and PDBeFold, while the SARS-CoV-2 similarity was also detected by HMI-PRED (Fuentes-Prior et al., 2000). Studies have shown that both thrombin and thrombomodulin blood concentrations are correlated with SARS-CoV-2 infection severity (Goshua et al., 2020; Ranucci et al., 2020). The verification by three tools and relevance to the literature led us to explore the potential mimicking of thrombomodulin binding to thrombin by the SARS-CoV-2 RBM using protein-protein docking (Figure 3A). Calculation of the interaction energies revealed that the reference docking and experimental controls showed similar affinities of -6.12 and -6.88 kJ/mol, and the SARS-CoV-2 RBM bound at a slightly lower affinity of -1.96 kJ/mol. The similarity to thrombomodulin might help explain the prothrombotic coagulopathy presented in SARS-CoV-2 infections (Bongiovanni et al., 2021).

**Figure 3.**
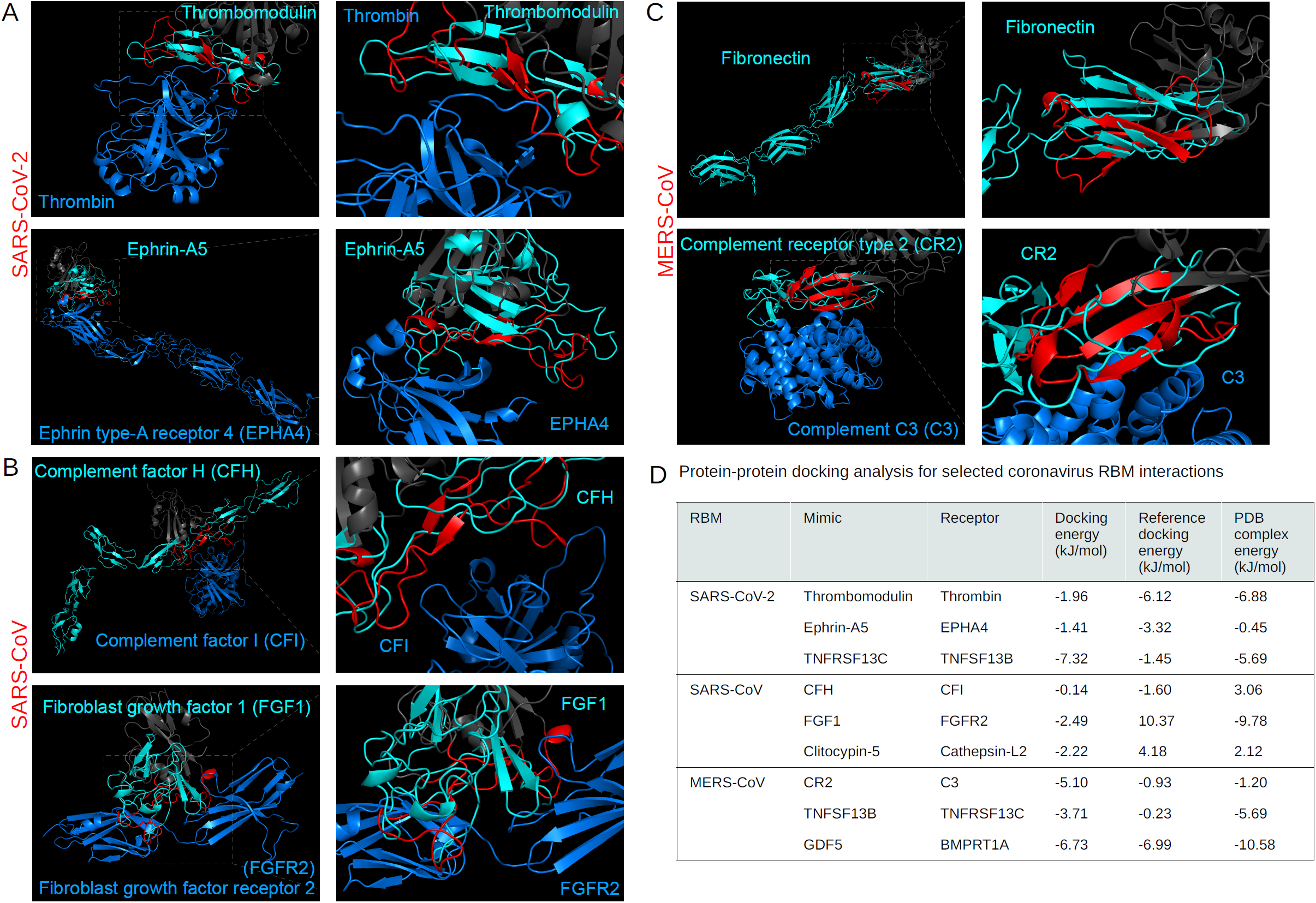
Analysis of endogenous structural alignments. Protein-protein docking was performed using ClusPro PIPER to test the potential interactions between the coronavirus RBMs and potential interaction partners. The following alignments are shown: between the SARS-CoV-2 RBM and thrombin (A, top) and ephrin type-A receptor 4 (A, bottom), the SARS-CoV RBM and complement factor I (B, top) and fibroblast growth factor receptor 2 (B, bottom), and the MERS-CoV RBM with complement C3 (C, bottom). The MERS-CoV RBM aligned to fibronectin type III domain (C, top). RBMs are labelled red, the remainder of the RBDs are dark gray, mimicked proteins are cyan, and potential interaction partners are marine blue. Interaction energy scores predicted using FoldX on docked and experimental complexes (D).

The central beta-strands of the SARS-CoV-2 RBM were found to be structurally similar to the transforming growth factor alpha, epiregulin, and epigen EGF-like domains using RUPEE; however, alignment of the RBM with the proteins in complex with the epidermal growth factor receptor (EGFR) (PDBs: 1mox, 5wb7, 5wb8) showed that the RBM was just out of ligand-binding range (Supplementary Figure 2E) (Freed et al., 2017; Garrett et al., 2002). There is no evidence for interaction of the SARS-CoV-2 RBM with the extracellular domain of EGFR; however, the alignments were included for potential off-target effects related to the EGF-like domains.

Structural mimicry of chemokine and cytokine signaling has been reported for several viruses (Alcami, 2003). Viral proteins can mimic the chemokine, as in the case of HIV gp120 and CCL5, or they can mimic the receptor and bind directly to the cytokine (inhibiting its function), such as the vaccinia virus B15R protein that mimics the IL-1B receptor and binds to IL-1B (Ahuja and Murphy, 1999; Alcami and Smith, 1992).

Several cell signaling ligands and receptors were found similar to the coronavirus RBMs. The SARS-CoV-2 and SARS-CoV RBMs were both found to mimic IL-8 like chemokines, fibroblast growth factor 1, C-C motif chemokines 3, interleukin-18, and ephrins; they also individually mimic BMP2 and von Willebrand factor, respectively. The MERS-CoV RBM structurally resembled C-X-C motif chemokines 2 and 4 and growth/differentiation factor 5. The alignments of the RBMs with IL-8 like chemokines, C-C motif chemokines 2 (CCL2), 3, and 4, and IL-18 in complex with their respective receptors shows only partial alignment with the ligand-binding regions (Supplementary Figure 2F). Interestingly, however, expression levels of these cytokines have all been shown as correlating with SARS-CoV-2 infection, although other explanations have been proposed (Buszko et al., 2020; Chen et al., 2010a; Jamilloux et al., 2020; Younes et al., 2020). For example, IL-33 release by damaged lower respiratory cells during SARS-CoV-2 has been demonstrated to trigger inflammation, increasing CCL2 and CCL3 expression (Zizzo and Cohen, 2020).

Fibroblast growth factor 1 (FGF1) was shown to be similar to the SARS-CoV-2 and SARS-CoV RBMs using PDBeFold and HMI-PRED. A transcriptomic profiling revealed that FGF1 was upregulated in coronavirus infections (Alsamman and Zayed, 2020). Thus, to look more closely at potential interference, we docked the SARS-CoV RBM with the fibroblast growth factor receptor 2 (FGFR2) (PDB: 3OJ2), which was predicted by HMI-PRED (Figure 3B) (Beenken et al., 2012). The RBM-FGFR2 docking analysis predicted a potentially favourable affinity of -2.49 kJ/mol, although not as high as the experimental complex (−9.78 kJ/mol) (Figure 3D). The FGF1 signaling pathway may, thus, be modulated by the coronavirus spike RBMs.

HMI-PRED predicted that the SARS-CoV-2 RBM mimics ephrin-A5 and ephrin-B2 binding to the ephrin type 4a receptor (EPHA4), and SARS-CoV mimics ephrin-A5 binding to the ephrin receptor type 3a. EPHA4 is unique among known class A ephrin receptors in that it binds both ephrin a and b ligands (Bowden et al., 2009; Xu et al., 2013). The structural similarity of two ligands for the same receptor for the SARS-CoV-2 RBM motivated further testing with protein-protein docking with EPHA4. Although there is no evidence for ephrin receptor involvement in coronavirus infections, other viral surface proteins have been shown to utilize ephrin receptors for cell entry, such as the rhesus r virus (Wang et al., 2020). The docking revealed similar affinities between the SARS-CoV-2 RBM-EPHA4, ephrin-A5-EPHA4, and experimental (PDB: 4m4r) complexes: -1.41, -3.32, -0.45 kJ/mol, respectively (Figure 3A).

The platelet glycoprotein Ib (GP-Ib) binding domain of von Willebrand factor (VWF) was found to be similar to SARS-CoV by HMI-PRED. VWF-GP-Ib interaction has been shown as critical in modulating thrombosis and inflammation (Denorme et al., 2019). Although there is no literature on VWF and SARS-CoV infection, blood concentration levels of VWF have been shown as correlated with SARS-CoV-2 infection severity, which may indicate potential pathway interference (Klok et al., 2020).

SARS-CoV-2 was predicted to mimic bone morphogenetic protein 2 binding to activin receptor type-2B and MERS-CoV to mimic growth/differentiation factor 5 (GDF5) binding to bone morphogenetic protein receptor type-1A (BMPRT1A) by HMI-PRED; however, no experimental evidence is available for either case. To explore the potential involvement of the MERS-CoV RBM in cell signaling, we docked the MERS-CoV RBM to the BMPRT1A in the GDF5-binding region (PDB: 3qb4) (Klammert et al., 2015). The docking of the RBM and GDF5 displayed similar affinities to BMPRT1A with -6.73 and -6.99 kJ/mol, respectively, while the experimental complex bound with -10.58 kJ/mol.

RUPEE detected structural resemblance between the SARS-CoV-2 RBM and IL-6 receptor alpha and beta chains, both of which show mimicry of the IL-6 binding sites (Supplementary Figure 2G). IL-6 has been reported as an overexpressed cytokine in SARS-CoV-2 infections, which can lead to induction of a hyper-innate inflammatory response (Chen et al., 2020; Magro, 2020). Mimicry of the IL-6 receptors by the RBM could result in binding and, thus, interference of IL-6 related interactions. However, several alternative theories have been proposed to explain the increases in IL-6 during severe infection; for example, the SARS-CoV nucleocapsid protein has been shown to activate IL-6 expression (Zhang et al., 2007). HMI-PRED additionally predicted MERS-CoV RBM mimicry of the binding of the T cell receptor beta chain to the major histocompatibility complex class I-related gene protein and interferon lambda receptor 1 binding to the beta subunit of the interleukin-10 receptor, both of which could have implications in immunosurveillance and inflammatory pathways (Corbett et al., 2014; Mendoza et al., 2017).

Different tumor necrosis factor-related ligands and receptors were found to be structurally analogous to the MERS-CoV RBM and SARS-CoV-2 RBM, respectively. Tumor necrosis factor receptor superfamily (TNFRSF) 1A, 4, 13C, and 14 were returned for the cystine disulfide loop for the SARS-CoV-2 RBM by RUPEE, while the tumor necrosis factor ligand superfamily (TNFSF) 13B and 14 were found to resemble the MERS-CoV RBM by RUPEE and HMI-PRED. Similarity of the SARS-CoV-2 RBM to TNFRSF 13C was also found by PDBeFold. These signaling pathways normally promote B-cell and the T-cell survival and maturation (Tamada et al., 2000; Yu et al., 2000). The structural similarity of this family of ligands and receptors to the SARS-CoV-2 and MERS-CoV RBMs led us to further inspect the interactions with protein-protein docking: mimicry of SARS-CoV-2 to TNFRSF 13C and MERS-CoV to TNFSF 13B. Thus, we simulated the binding of the SARS-CoV-2 RBM to TNFSF 13B and MERS-CoV to TNFRSF 13C (PDB: 3v56) (Smart et al., 2012). Both cases revealed that the RBM is predicted to dock at a higher affinity than the natural ligand (Figure 3D).

The complement system comprises a series of protein cascades that form an integral part of the innate immune response to viruses (Nonaka and Yoshizaki, 2004). Viruses are generally susceptible to the complement system; however, viral proteins can utilize complement proteins through molecular mimicry in a variety of ways, such as using complement receptors for viral entry or evading detection by the immune system (Bernet et al., 2003). Infections from all three highly pathogenic coronaviruses have been reported to activate the complement system, enhancing pathogenicity, although the exact mechanisms remain unclear (Java et al., 2020). The spike protein of SARS-CoV-2 has been shown to localize near C4d and C5b-9 in lung vasculature, and mutations in several complement activation proteins, such as complement factors H, I, and III, have been found to correlate with infection severity (Magro et al., 2020; Ramlall et al., 2020). The structural similarity screen yielded three motifs from the complement system that potentially mimic RBMs: complement factor I (CFI) binding domain of CFH for all three RBMs and both the complement C3d binding domain of complement receptor 2 (CR2) and the complement C1r binding domain of complement C1s for the MERS-CoV RBM. Interestingly, CFH and the SARS-CoV-2 spike protein have been proposed to compete for heparan sulfate binding (Yu et al., 2020). The SARS-CoV RBM, however, was predicted to be similar to CFH by RUPEE, PDBeFold, and HMI-PRED; thus, we docked the SARS-CoV RBM to CFI (PDB: 5o32) and found that the natural ligand was predicted to bind at a slightly higher affinity than the SARS-CoV RBM: -1.60 vs. -0.14 kJ/mol, respectively (Xue et al., 2017). The C3d-binding domain of CR2 for the MERS-CoV RBM was also identified by RUPEE, PDBeFold, and HMI-PRED and was, thus, explored with docking of the MERS-CoV RBM to C3 (PDB: 3oed) (Figure 3C) (van den Elsen and Isenman, 2011). The MERS-CoV RBM was predicted to bind at a higher affinity than both the control docking and experimental complexes: -5.10 vs. -0.93 and -1.20 kJ/mol, respectively (Figure 3D). Additionally, HMI-PRED found that the MERS-CoV RBM also mimics the complement C1r binding site of complement C1s. Additional experimental efforts are needed to validate the relationship between coronavirus spike proteins and the complement activation pathway.

Other endogenous hits included several unrelated proteins, such as protease inhibitors and serotransferrin. The MERS-CoV RBM resembled the fibronectin type III (FNIII) domains of mouse myosin-binding protein C and tenascin-X using RUPEE. Although myosin-binding protein C is intracellular, FNIII domains are found across the domains of life and function in diverse ways, from cell adhesion to cell signaling (Campbell and Spitzfaden, 1994). Drayman *et al*. found that the West Nile virus envelope glycoprotein E resembles the structural architecture of the FN10 domain of fibronectin, which is a natural ligand for integrin αvβ3. Thus, we checked and found that the MERS-CoV RBM shares structural properties with other FNIII domains, such as those from fibronectin and neural cell adhesion molecule 1 (PDBs: 2haz and 1fnf, respectively) (Figure 3C) (Leahy et al., 1996; Mendiratta et al., 2006). The MERS-CoV RBM was also found to mimic part of the jagged-2, DLL1, and DLL4 proteins; however, the alignment was largely out of ligand-binding range when compared to jagged-1 in complex with NOTCH1 (PDB: 5uk5) – although the alignment may be relevant in other scenarios (Supplementary figure 2E) (Luca et al., 2017). Protease inhibitors included neuroserpin for the SARS-CoV-2 RBM and clitocypin-5 cysteine protease for the SARS-CoV RBM. The alignment of the SARS-CoV with clitocypin-5 cysteine protease showed potential binding to cathepsin L2 (PDB: 3h6s) (Renko et al., 2010). The role of cathepsins in coronavirus cell entry has been described as helping process the spike protein for viral and host membrane fusion (Pišlar et al., 2020). To investigate the potential for additional interactions between coronavirus RBMs and cathepsins, we performed protein-protein docking. The binding of the SARS-CoV RBM to cathepsin L2 was predicted to be more favourable than the docking and experimental controls (Figure 3D). Experimental evidence is required to validate this interaction, however. Both the SARS-related RBMs resembled motifs of serotransferrin using TM-align, and, interestingly, the transferrin receptor protein 1 has been proposed as a potential cell entry receptor for SARS-CoV-2 (Tang et al., 2020). However, the alignments were generally out of ligand-binding range (PDB: 1suv) (Supplementary Figure 2H); since no binding mode was apparent, it was not considered for docking (Cheng et al., 2004). HMI-PRED predicted that the SARS-CoV-2 RBM mimics the dimerization domain of C-type lectin domain family 5 and that the SARS-CoV RBM mimics intercellular adhesion molecule 5 binding to integrin alpha-L (Watson et al., 2011; Zhang et al., 2008). Integrins have been proposed to bind to the SARS-CoV-2 spike protein, although that is due to a new RGD motif in the RBD – of note, the RGD motif is not included in the selected residues for this study’s SARS-CoV-2 RBM since it does not interact with ACE2 in experimental models (Sigrist et al., 2020). Because integrin binding has not been hypothesized outside of the discussion of the SARS-CoV-2 RGD motif, docking was not pursued. Both the SARS-CoV-2 and SARS-CoV RBMs mimicked the nicotine-binding domain of the nicotinic acetylcholine receptor by RUPEE, which may have implications in the ‘nicotinic hypothesis’ (Changeux et al., 2020).

#### Exogenous

We classified the exogenous hits by the pathogen type. There were motifs from apicomplexan parasites, viruses, one bacterial protein, and snake and spider toxins found to resemble the coronavirus RBMs.

The EGF-like domains from merozoite surface protein 1 (MSP1) of several *Plasmodium* species were found to be structurally similar to all three RBMs using RUPEE. Compared to the other two, the SARS-CoV-2 RBM was found to be similar to the most *Plasmodium* species: *falciparum, yoelii, cynomolgi, knowlesi, vivax*. The SARS-CoV RBM returned *P. yoelii* MSP1 and the MERS-CoV RBM returned *P. falciparum* MSP1. A closer inspection at the *P*. f*alciparum* MSP1 alignments revealed that two EGF-like domains on the same PDB structure (1ob1) were found to resemble the SARS-CoV-2 RBM (Figure 4A) (Pizarro et al., 2003). The PDB structure is originally modelling the antibody-binding epitope of the EGF-like domain of MSP1; however, the antibody epitope is located on a loop just outside of the EGF-like domain. Thus, antibody-binding to the SARS-CoV-2 RBM could not be verified, but the presence of two EGF-like domains near an epitope may motivate experimental testing. The *P*. f*alciparum* apical membrane antigen 1 epitope (PDB: 2j5l) was also found to resemble the SARS-CoV-2 and SARS-CoV RBMs, although, as in the case of MSP1, both RBMs aligned to a region outside of the antibody-interacting residues (Figure 4B) (Igonet et al., 2007). These EGF-like domains from *Plasmodium* parasites may provide structural epitope scaffolding for cross-reactivity against the coronavirus spike RBMs (Craig et al., 1998). Recent studies have pointed to a potential protective effect of *P. falciparum* infections against SARS-CoV-2 infection, although direct experimental evidence is yet to be established (Iesa et al., 2020; Kalungi et al., 2021; Panda et al., 2020; Raham, 2021). The MERS-CoV RBM was also found to resemble the rhoptry neck protein 2 and thrombospondin-related anonymous protein from *P. falciparum*. The surface antigen 3 of *Toxoplasma gondii* was found to be similar to the MERS-CoV RBM (Figure 4C). Although there are no data on MERS-CoV and *T. gondii* co-infections, SARS-CoV-2 has been shown to have negative covariation with toxoplasmosis, which may indicate a protective effect from *T. gondii* (Jankowiak et al., 2020).

**Figure 4.**
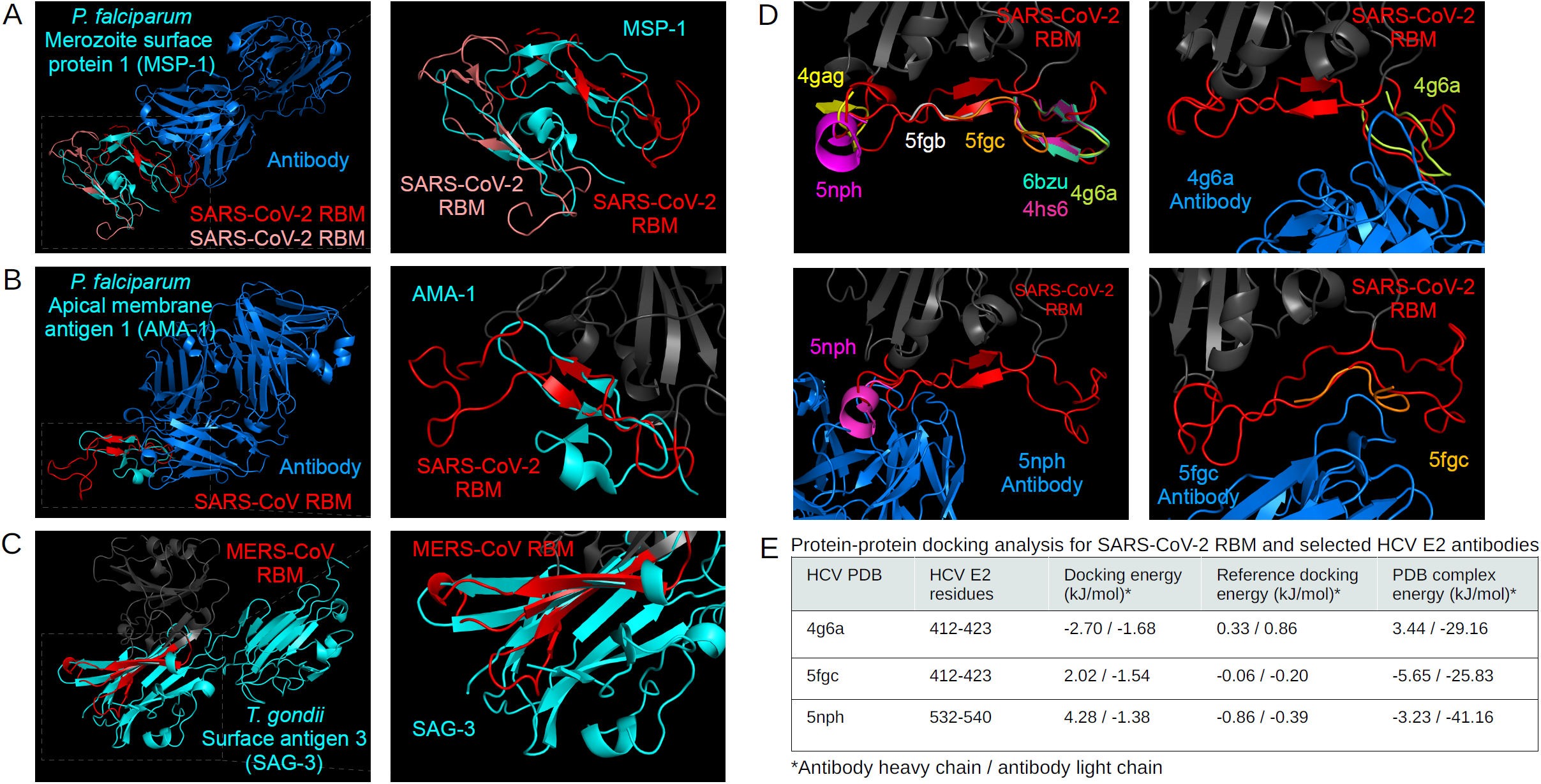
Analysis of exogenous structural alignments. *Plasmodium falciparum* merozoite surface protein 1 (A) and apical membrane antigen 1 (B) structurally aligned with the SARS-CoV-2 RBM and SARS-CoV RBM, respectively. The *Toxoplasma gondii* surface antigen 3 aligned with the MERS-CoV RBM (C). RBMs are labelled red, mimicked proteins are cyan, and potential interaction partners are marine blue (A,B,C). Hepatitis C virus epitopes structurally aligned to the SARS-CoV-2 RBM, and the respective antibody structures from PDBs 5fgc, 5nph, 4g6a docked to the RBM (D) using ClusPro PIPER “antibody” mode. Interaction energy scores predicted using FoldX on docked and experimental complexes (E).

The coronavirus RBMs were found to structurally mimic several motifs on the HIV and Influenza spike proteins; however, they were found either facing inwards or buried inside the mimicked protein and were, therefore, discarded. PDBeFold and TM-align indicated that the SARS-CoV-2 and SARS-CoV RBMs structurally mimic several hepatitis C virus (HCV) antibody epitopes. The SARS-CoV-2 and SARS-CoV RBMs were found to be similar to 10 and 6 PDB HCV E2 protein epitopes structures, respectively (Supplementary Table 2). The HCV E2 protein is implicated in host entry, which has been explored as an inhibitory target with neutralizing antibodies (Heile et al., 2000; Ploss and Evans, 2012). A closer inspection of the mapping of the epitopes to the SARS-CoV-2 RBM show that they are distributed across the RBM (Figure 4D). Some studies have suggested that HCV may be negatively correlated with SARS-CoV-2 infection (Mirzaie et al., 2020; Reddy, 2020; Richardson et al., 2020). Since several of the epitopes were aligned in ways that were accessible to antibodies in the original PBD, we selected three epitopes, one at each region of the RBM, and docked the respective antibody to the SARS-CoV-2 RBM using the ClusPro PIPER ‘antibody’ mode (Figure 4D). As shown in Figure 4E, the RBM-antibody docking results were compared to docking and experimental controls – the antibodies bound in a similar way to docking controls in all three cases, while the experimental complexes were predicted to bind more tightly. These structural similarities may take part in potential cross-reactivity between HCV and coronavirus infections. Of note, two recently proposed cell entry receptors for the SARS-CoV-2 spike protein, ASGR1 and APOA4, have been shown as potentially implicated in mediating HCV viral entry (Brockbank et al., 2021). In an interesting case, the MERS-CoV RBM was found to structurally mimic both the Machupo virus glycoprotein polyprotein GP complex RBM (TM-score: 0.47) and its receptor, transferrin receptor protein 1 (0.52) (PDB: 3kas) using TM-align, although the transferrin receptor scores slightly higher (Supplementary Figure 3A) (Abraham et al., 2010).

Only one bacterial protein was selected in the structural similarity screen. The adhesin-binding fucosylated histo-blood group antigen of *Helicobacter pylori* was found to be similar to the MERS-CoV RBM by TM-align. The structure (PDB: 5f7l) shows binding of the bacterial protein to a nanobody; however, the RBM alignment is just outside of the nanobody binding site (Supplementary Figure 3B) (Moonens et al., 2016). No studies have detailed any connections between MERS-CoV and *H. pylori*.

Motifs from snake, spider, and cone snail toxins were found to be similar to all three RBMs using PDBeFold and RUPEE. The SARS-CoV-2 and SARS-CoV RBMs shared similarity to four toxins, and the MERS-CoV RBM only returned unique proteins. The two SARS-related viruses mimicked three-finger bungarotoxins and inhibitor cystine-knot toxins, such as psalmotoxin-1, while MERS-CoV RBM resembled other three-finger toxins, like cytotoxin 4 (Supplementary Figures 3C,3D) (Corfield et al., 1989; Dellisanti et al., 2007; Lee et al., 2014; Saez et al., 2011; Volpon et al., 2004). In total, these toxins may bind to several receptors involved in nociception, e.g. ASIC1 and Nav1.7, which may be relevant to the taste and pain perception changes experienced during SARS-CoV-2 infection (McFarland et al., 2021). Importantly, and perhaps confoundingly, a recent study found no changes in depolarization for Nav1.7 and Cav2.2 upon exposure to the SARS-CoV-2 RBD (Moutal et al., 2020). Thus, further experimental work is necessary to validate these interactions.

## Conclusions

This study involved the structural bioinformatics characterization of potential molecular mimicry by highly pathogenic coronavirus spike protein RBMs. Using protein homology modelling, we built representative models of the spike RBMs and tested structural changes in the SARS-CoV-2 RBM induced by recently recorded mutations, which had little effect on overall RBM structure. Comparison of the RBMs revealed that the SARS-CoV and SARS-CoV-2 RBMs share higher structural homology than with MERS-CoV, which was underlined by the number of common returned proteins in the structural similarity screen using four structural alignment tools. The flexibility of the cystine disulfide loop in the SARS-related RBMs was found to permit large global changes in RBM structure; however, since most of the predicted mimicry was mapped to the RBM central beta-strands, which are quite rigid, the models of different conformations did not return significantly different proteins from the structural alignment tools. The structural alignment screen highlighted the similarity of the RBMs to evolutionarily unrelated human and non-human proteins. Further validation of the alignments with protein-protein docking revealed that all tested coronavirus RBM-endogenous protein interactions were predicted to be energetically favourable, confirming that the structural similarity screen may be useful in identifying potential molecular mimics.

The predicted endogenous mimicry comprised of proteins in cell signaling, adhesion, and complement pathways. Potential mimicry of several microbial antigenic proteins and exogenous toxins was also discovered. The EGF-like domains of both endogenous and exogenous proteins structurally resemble all three RBMs. Predicted mimicked endogenous interactions include the EGF-like domain of thrombomodulin binding to thrombin, NOTCH1 binding to DLL4, and coagulation factor VIIa binding to tissue factor. Interference in these pathways may partially explain coagulopathies in coronavirus infections (Iba et al., 2020). Exogenous EGF-like domains of MSP1 from different *Plasmodium* species, on the other hand, may provide a structural epitope scaffold for cross-reactivity between coronavirus and *Plasmodium* infections (Panda et al., 2020). Epitope similarity was further explored among the several antibody-bound hepatitis C virus E2 protein motifs that were structurally analogous to the SARS-related RBMs. Structural similarity to antigenic proteins from other microbes may confer cross-immunity and, thus, also potentially guide vaccine design. Cell signaling pathway proteins, such as TNF-related and ephrin ligands, were also found as potential mimics of the coronavirus RBMs, which may lead to use of alternative co-receptors for viral entry or modulation of signaling cascades. Complement factor H was returned for all three RBMs and has also been implicated in coronavirus infections (Yu et al., 2020). The mimicry of complement proteins is widespread among viruses, and the spike RBM may have secondary roles interfering in these pathways (Mastellos et al., 2003). Many snake and spider toxins were also found similar to the coronavirus RBMs, which implies the potential usage of receptors involved in pain, muscle contraction, cell adhesion, and coagulation pathways (Dongol et al., 2019; Rowan, 2001; Wu et al., 2006). The prediction of evolutionarily unrelated, yet structurally similar, potential protein mimics reveals that previously unidentified pathways could be altered by the spike RBMs. The structural variation between coronavirus RBMs and their resulting molecular mimics can possibly be connected to differences in tropism, infection severity, and immune system reactivity between coronaviruses.

Although experimental verification of the predicted interactions is required to take these results further, the findings presented in this study provide insight into the potential molecular mimicry utilized by highly pathogenic coronavirus RBMs. The data can be used to support inhibitory drug, peptide, and antibody design efforts in order to prevent viral cell entry and virulence mechanisms related to coronavirus RBMs. Additional work is needed to better understand how coronaviruses co-opt host machinery to enhance fitness.

## Supporting information

Manuscript_tables

## Acknowledgements

Many thanks to the ClusPro PIPER team for letting our team download and install their software. A big thank you to the BioRender team and Melissa Pappas for help with figure rendering. CAB, SCV, and TLB were supported by American Leprosy Missions, United States of America (Grant No: G88726). TLB thanks the Wellcome Trust for support through an Investigator Award (200814/Z/16/Z; 2016 -2021). ARJ is funded by the Biotechnology and Biological Sciences Research Council (BBSRC) DTP studentship (BB/M011194/1). AFA is funded by the King Abdullah scholarship (Saudi Arabia Research Council). LC and TLB are being funded by Ipsen Bioinnovation (PHZJ/487 -RG83343). AvT was supported by a joint Biotechnology and Biological Sciences Research Council (BBSRC) research grant under grant number BB/N00468X/1. BPB is supported by The Wellcome Trust (107032AIA), The UK Cystic Fibrosis Trust (Innovation Hub grant 001) and NIHR Cambridge Biomedical Research Centre. SET is funded by the Cystic Fibrosis Trust (RG 70975) and Fondation Botnar (RG91317). PHMT is funded by the Brazilian National Council for Scientific and Technological Development (CNPq).

## Author contributions

CAB performed analyses and wrote the paper; ARJ, AFA, and LC performed analyses and provided expertise on bioinformatics tools and statistical analysis; AvT, SH, BPB, and SET provided expertise on sequence and structural investigations; SCV helped acquire funding and provided expertise on the structural mutation analysis; PHMT and TLB supervised and provided overall direction for the study.

## Declaration of interests

The authors declare no competing interests.

## Methods

### Spike receptor-binding motif model generation and characterization

Amino acid sequences of the SARS-CoV-2 (NCBI code: NC_045512), SARS-CoV (NC_004718), and MERS-CoV (NC_038294) spike proteins were extracted as FASTA files from the NCBI Viral Genomes Resource (Brister et al., 2015). Each amino acid sequence corresponds to one of three identical protomers of the full homo-oligomeric spike trimer. Due to the high number of available experimentally-resolved structures for each spike protein, representative models were generated using ProtCHOIR – a recently developed bioinformatic tool to automate 3D homology modelling of homo-oligomers (Torres and Blundell, manuscript in preparation https://doi.org/10.5281/zenodo.3384945). ProtCHOIR builds homo-oligomeric assemblies by searching for homolog templates on a locally created homo-oligomeric protein database using PSI-BLAST, performing a series of structural analyses on the input protomer structure or sequence using Molprobity, PISA, and GESAMT (all three tools as part of the CCP4 Molecular Graphics package), and comparative homology modelling using MODELLER (version 9.24) with molecular dynamics-level optimization and refinement (Altschul et al., 1997; Chen et al., 2010b; Krissinel, 2012; Krissinel and Henrick, 2007; Šali, 2019; Šali and Blundell, 1993).

The residues for the receptor-binding domains (RBD) and RBMs of each spike model were manually selected based on experimental structures with primary receptors (residues defined in Supplementary Table 1) and made into sub-structures during manual inspection of full-length models on PyMOL (The PyMOL Molecular Graphics System, Version 2.0 Schrödinger, LLC). Global amino acid sequence alignment of RBDs was performed with EMBOSS Needle (Madeira et al., 2019). The full-length SARS-CoV-2 spike protein modelled with the lipid bilayer displayed in Figure 1 was retrieved from the SARS-CoV-2 3D database (Alsulami et al., 2021). No records exist of N-linked or O-linked glycosylation motifs near the three RBMs, which was supported by NetNGlyc 1.0 and NetOGlyc 4.0 predictions (Gupta et al., 2004; Steentoft et al., 2013). To determine the flexibility of each residue in the RBDs, we used CABS-flex 2.0, a web server that offers fast simulations and resulting data of protein structure flexibility (Kuriata et al., 2018). Default values were used and residue flexibility was reported as root mean squared flexibility (RMSF) (Kmiecik et al., 2016).

### Structure similarity screen

Several web servers and stand-alone tools have become available to perform pairwise or multiple sequence-independent protein structure alignments, such as DALI, FATCAT, iSARST, MADOKA, PDBeFold, TM-align, and RUPEE (Ayoub and Lee, 2019; Deng et al., 2019; Holm and Laakso, 2016; Krissinel and Henrick, 2004; Li et al., 2020; Lo et al., 2009; Zhang and Skolnick, 2005). After testing each tool, the PDBeFold web server, RUPEE web server, and a locally-installed version of TM-align were selected due to the diversity of structural alignment methodologies, ease-of-use, data accessibility, and widespread-usage. A newly published web server for structural prediction of host-microbe interactions based on interface mimicry, HMI-PRED, was also included in the analysis.

Of note, mTM-align (the web server version of TM-align) was considered, but no non-spike proteins were shown – restricting the downstream analysis (Dong et al., 2018). Thus, all 3D models in the PDB database clustered at 100% sequence identity were downloaded, and TM-align was run in a pairwise manner, using GNU parallel, between each RBM model and each chain of every downloaded PDB file (O. Tange (2018): GNU Parallel, March 2018, https://doi.org/10.5281/zenodo.1146014.). TM-align works by, first, combining secondary structure similarity alignments, defined by DSSP (Define Secondary Structure of Proteins), and TM-score-based structural alignments (Kabsch and Sander, 1983; Zhang and Skolnick, 2004). A structure rotation matrix is applied to the alignments in order to maximize the TM-score, which was used to rank the alignments for each RBM.

The PDBeFold web server utilizes SSM, a graph-matching algorithm that superimposes *PROMOTIF*-defined secondary structures and, subsequently, maps backbone carbon atoms of, first, matched and, second, unmatched secondary structures (Hutchinson and Thornton, 1996; Krissinel and Henrick, 2004, 2005). The hits are ranked by their Q-Score, which is calculated to achieve a lower root mean squared deviation (RMSD) and an increased number of aligned residues. Since the highest percentage (%) of secondary structure matches for the SARS-CoV RBM was found at 67% (while SARS-CoV-2 and MERS-CoV returned hits with 100% secondary structure matches), we set the PDBeFold search parameters for 65% structural similarity at “highest precision” for each RBM.

The RUPEE web server performs structural similarity comparisons using a purely geometric approach: 1) a linear encoding of the protein structure is defined to identify separable regions of permissible torsion angles for DSSP secondary structure assignments; 2) the encoding is converted into a bag of features; 3) a protein structure indexing method is established using min-hashing and locality sensitive hashing; 4) the top 8,000 matches are sorted based on adjusted Jaccard similarity scores; and 5) if running in “Top-Aligned” mode (used in this analysis), the alignments are re-scored using TM-align (Ayoub and Lee, 2019). RUPEE allows the specific comparison of a query protein to the CATH, SCOP, PDB, and ECOD databases (Andreeva et al., 2020; Burley et al., 2019; Cheng et al., 2014; Sillitoe et al., 2019). Additional settings are also offered, such as the “contains” (finding query protein inside database protein) and “contained in” options (small protein motif detection in query protein) – both of which were used in this analysis.

HMI-PRED combines TM-align with NACCESS to search through template host protein-protein complexes (Guven-Maiorov et al., 2020; Hubbard and Thornton, 1993). The structural alignment model of the ligand and putative receptor are refined with RosettaDock to quantify electrochemical complementarity, which is not included in a strict structural alignment screen (Wang et al., 2007). Alignments for both the RBM and RBD of each of the three coronaviruses were collected from HMI-PRED.

Since the flexibility of the cystine disulfide loop on the SARS-CoV-2 and SARS-CoV RBMs may affect the global structure of the RBMs and, thus, search outcome, we used two additional models provided by CABS-flex 2.0 for both RBMs, making a total of three conformations for each SARS-CoV-2 and SARS-CoV. Results of the different conformations provided were pooled together for both SARS-related RBMs. The top-scoring alignments from all four tools for each RBM were matched with their corresponding PDB and, subsequently, UniProt accession code (Bateman, 2019). The UniProt accession codes were then compared across tools to identify shared top hits.

### Protein-protein docking and interaction energy prediction

A local installation of ClusPro PIPER (version 1.1.5) was used for protein-protein docking (Desta et al., 2020; Kozakov et al., 2013, 2017; Vajda et al., 2017). Annotations informing potential protein-protein interactions were obtained from the PDB, STRING, and UniProt databases (Szklarczyk et al., 2019). The ClusPro PIPER “antibody” docking mode was used to dock RBDs with the hepatitis C antibody PDB structures (PDBs: 4g6a, 5fgc, 5nph), and the “others” mode was used for all non-antibody docking. In order to minimize non-biologically relevant binding during the docking runs, residues outside of the RBM on the spike RBDs and outside of the ligand-binding region on the predicted interaction partners were masked. The docked models were minimized using CHARMM22. To gain better insight into the binding strength of the potential RBD-receptor complex interactions, we used the FoldX (version 4.0) AnalyseComplex program, which predicts the interaction energy by finding the difference in stability between the individual unfolded molecules and the overall complex (Schymkowitz et al., 2005). The original PDB ligands were also docked to the receptor in order to obtain a “Reference docking energy” when compared with the predicted RBD-receptor energy. The “PDB complex energy” was obtained from the original PDB containing the ligand and receptor to understand binding resolution of the experimental complex.

### Data analysis and visualization

A full representation of the pipeline and tools used can be found in Figure 5. Data were analyzed and plots were generated using R version 3.6.3 (2020-02-29). Protein structural alignments were visualized with PyMOL (version 1.8.4.0) (Stout, 2004). Pdb-tools was used to manipulate and organize PDB files (Rodrigues et al., 2018). The graphical abstract was adapted from the “SARS-CoV-2 Spike Protein Conformations” template on BioRender. Figure 5 was created on draw.io. Raw data and alignment models are made available at https://github.com/tlb-lab.

**Figure 5.**
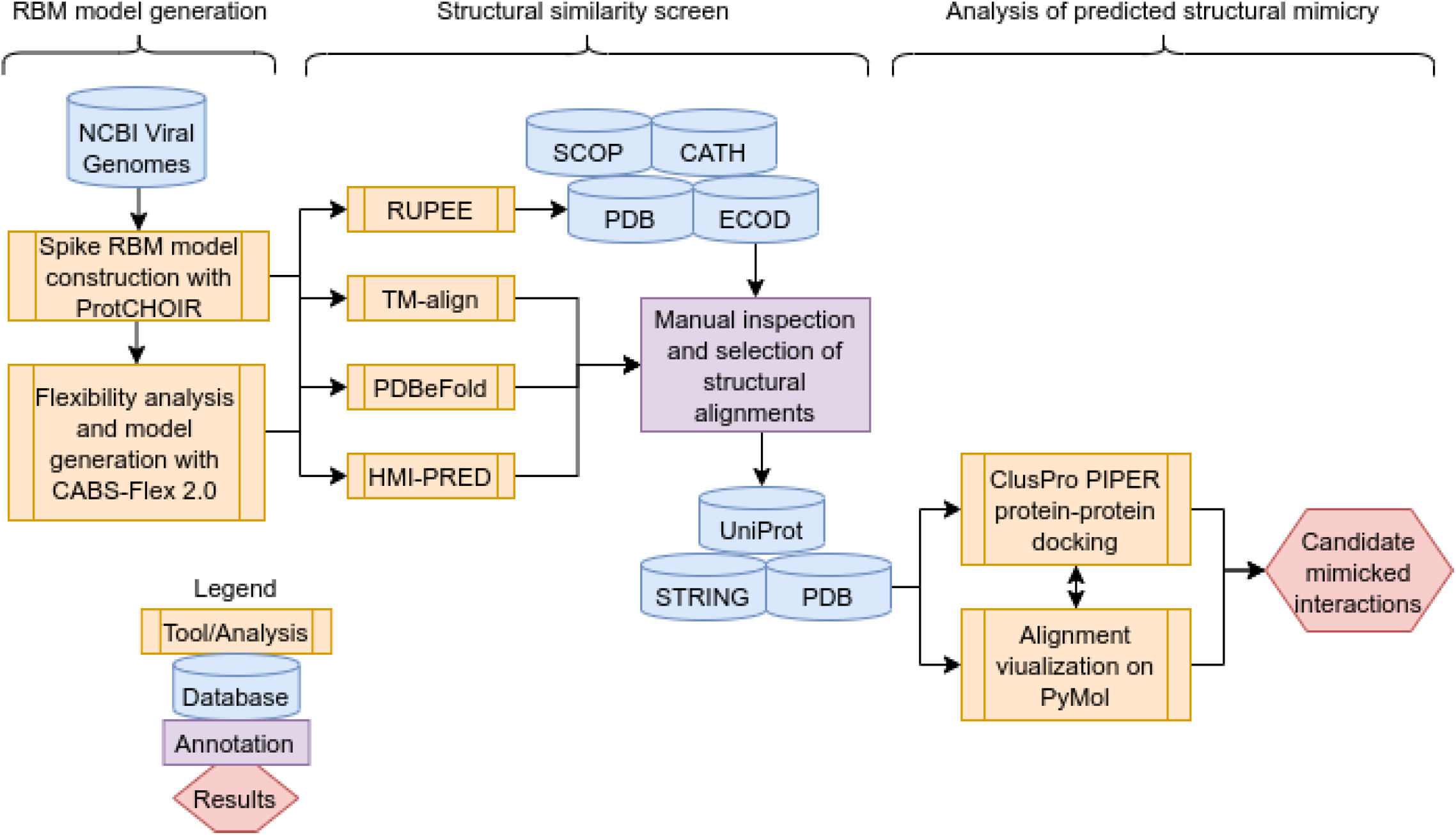
Pipeline flow. A flow chart of the analyses performed in this study.

## Supplementary figures and tables

**Supplementary Table 1.** Spike protein receptor-binding domain and motif residues selected in this study.

| RBM | Receptor-binding domain residues | Receptor-binding motif residues |
| --- | --- | --- |
| SARS-CoV-2 | 333-523 | 443-457,469-506 |
| SARS-CoV | 332-511 | 426-443,456-492 |
| MERS-CoV | 380-586 | 501-515,529-558 |

**Supplementary Figure 1.**
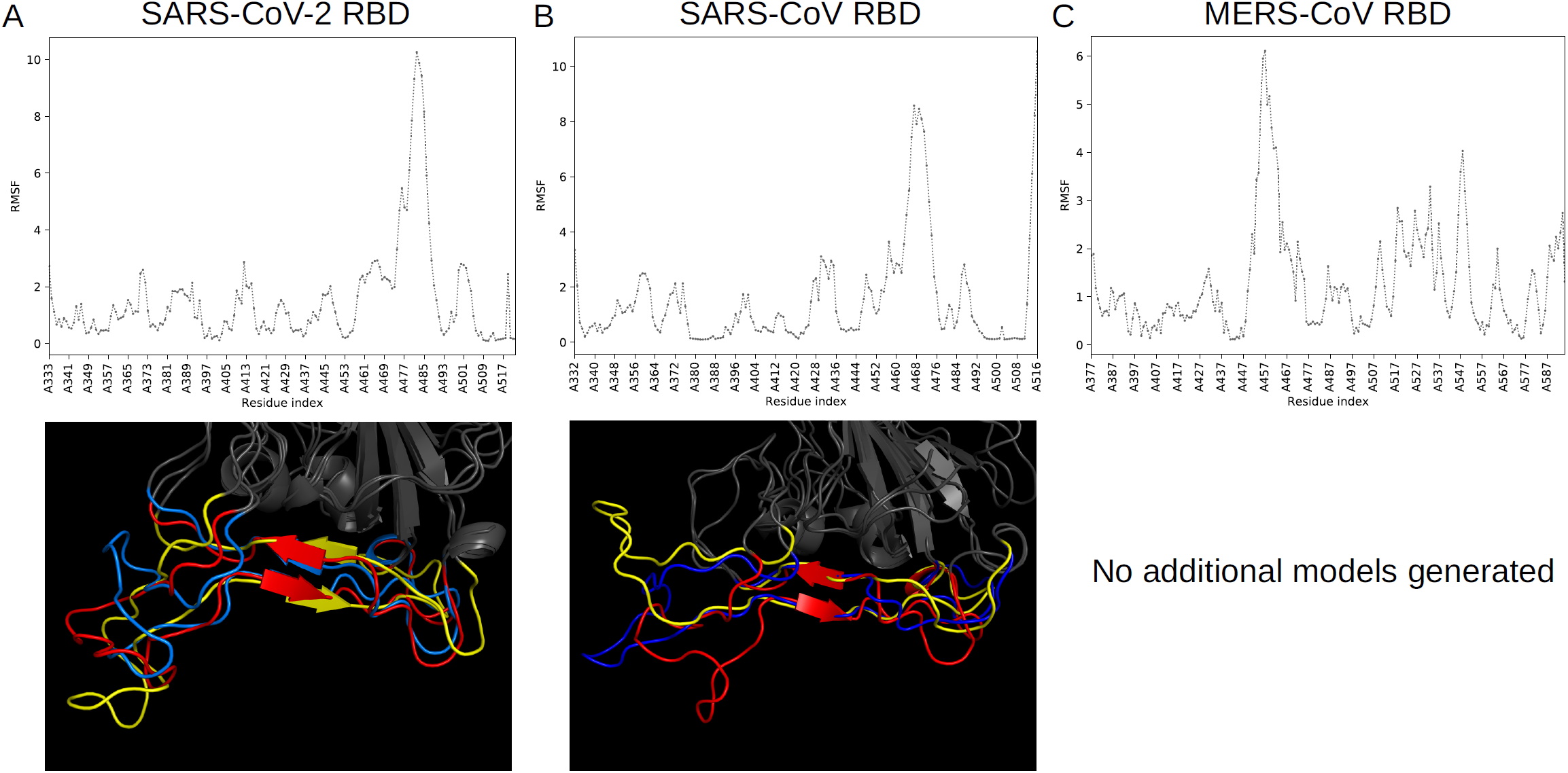
CABS-Flex 2.0 analysis and models for coronavirus RBDs. The top row displays the root mean square flexibility for each residue in the SARS-CoV-2 (A), SARS-CoV (B), and MERS-CoV (C) RBDs in the CABS-Flex 2.0 analyses. The bottom row shows selected models from CABS-Flex 2.0, included for the SARS-CoV-2 and SARS-CoV structural similarity screen, in alignment with the reference RBM.

**Supplementary Figure 2.**
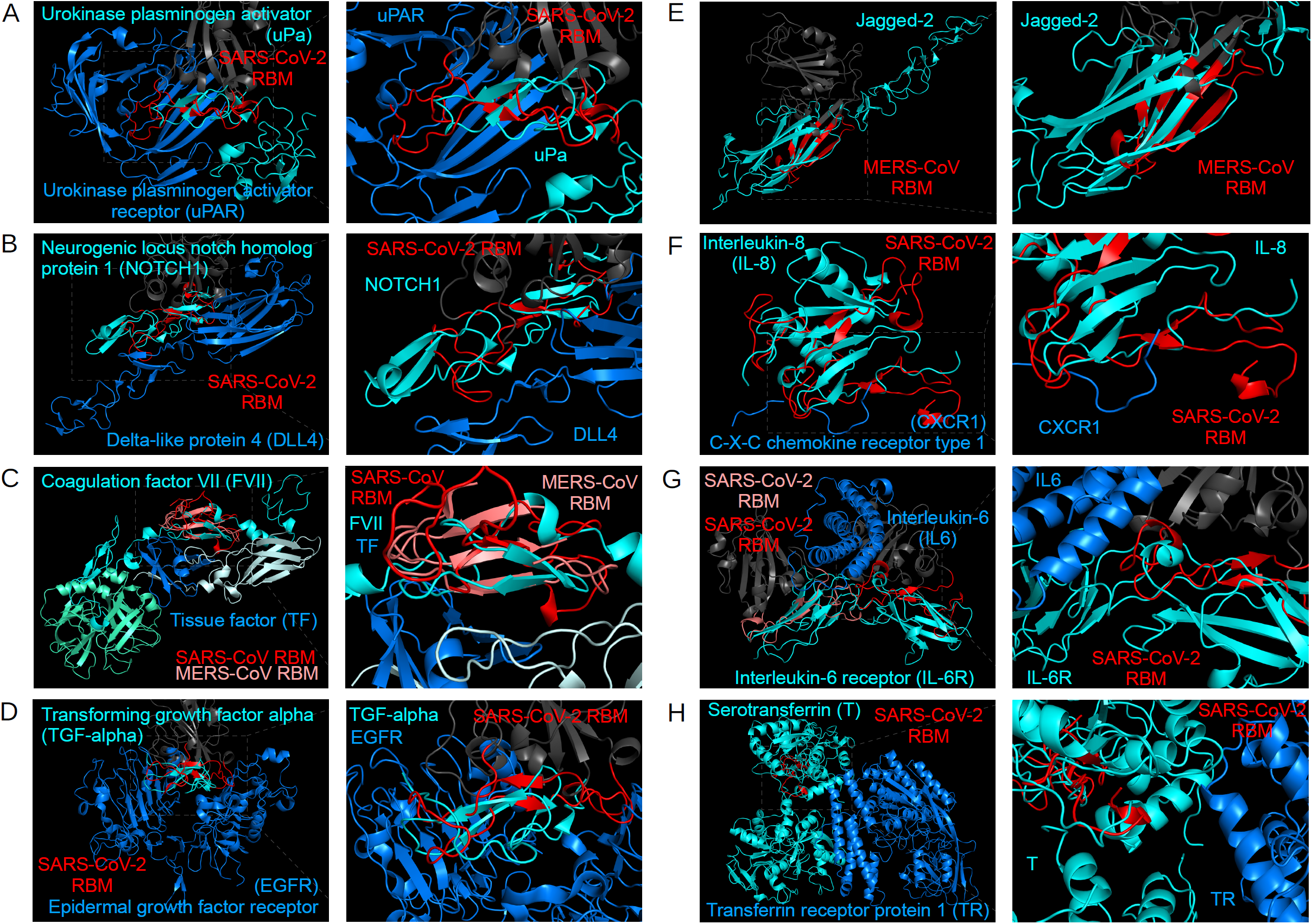
Structural alignments of potential endogenous mimics. Structural alignments between the following coronavirus RBMs and potential molecular mimics are shown: SARS-CoV-2 and urokinase plasminogen activator (A), SARS-CoV-2 and NOTCH1 (B), MERS-CoV and SARS-CoV mimicking coagulation factor VIIa (C), SARS-CoV-2 and TGF-alpha (D), MERS-CoV and jagged-2 (E), SARS-CoV-2 and IL-8 (F), SARS-CoV-2 and IL-6 receptor alpha and beta chains (G), SARS-CoV-2 and serotransferrin (H). RBMs are labelled red, mimicked proteins are cyan, and potential interaction partners are marine blue.

**Supplementary Figure 3.**
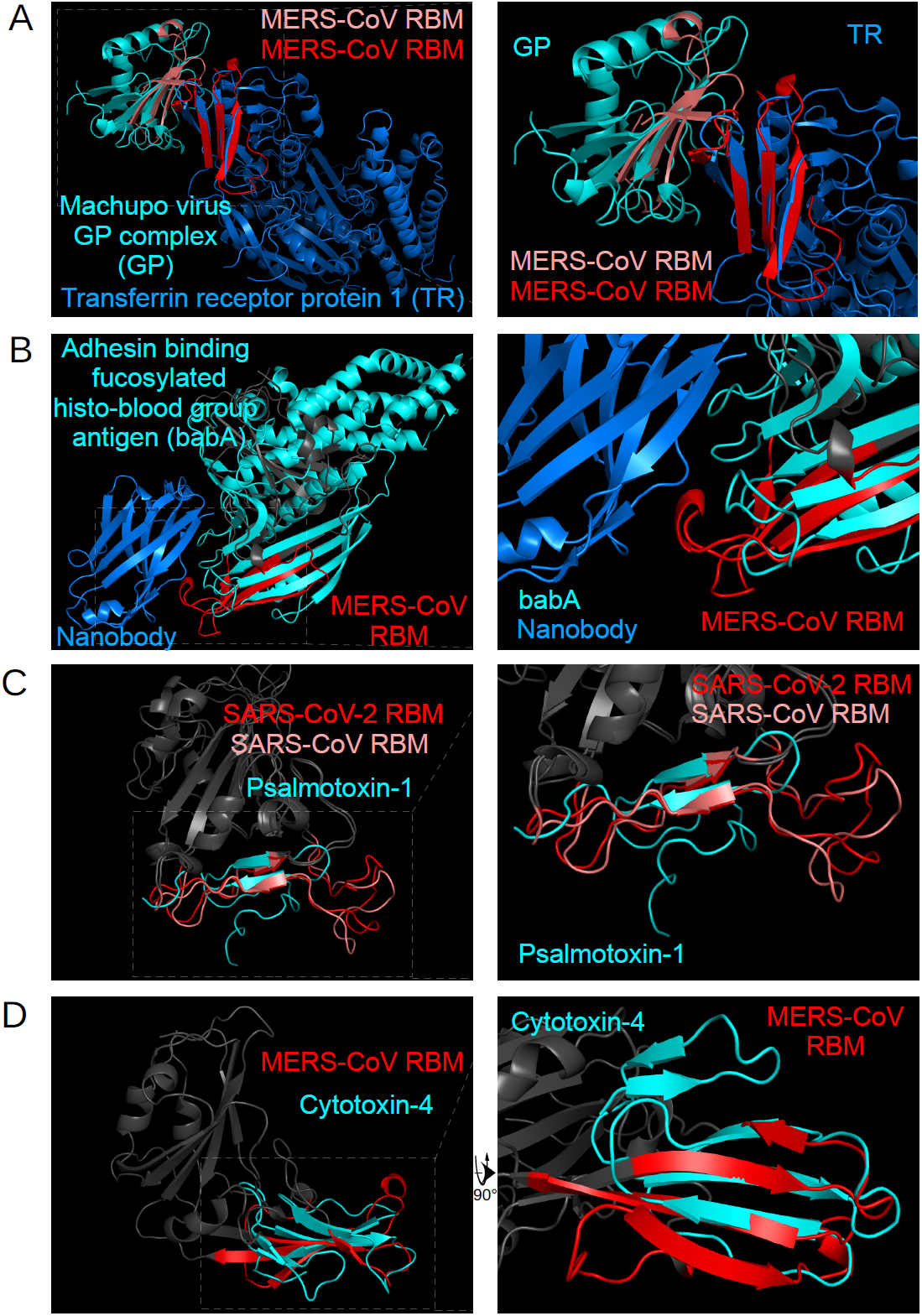
Structural alignments with exogenous proteins. The MERS-CoV RBM aligned to the transferrin receptor protein 1 and Machupo virus glycoprotein polyprotein GP complex (A) and the adhesin-binding fucosylated histo-blood group antigen (B). The SARS-CoV and SARS-CoV-2 RBMs aligned to psalmotoxin-1 (C), and the MERS-CoV RBM aligned to cytotoxin 4 (D). RBMs are labelled red, mimicked proteins are cyan, and potential interaction partners are marine blue.

**Supplementary Table 2.**
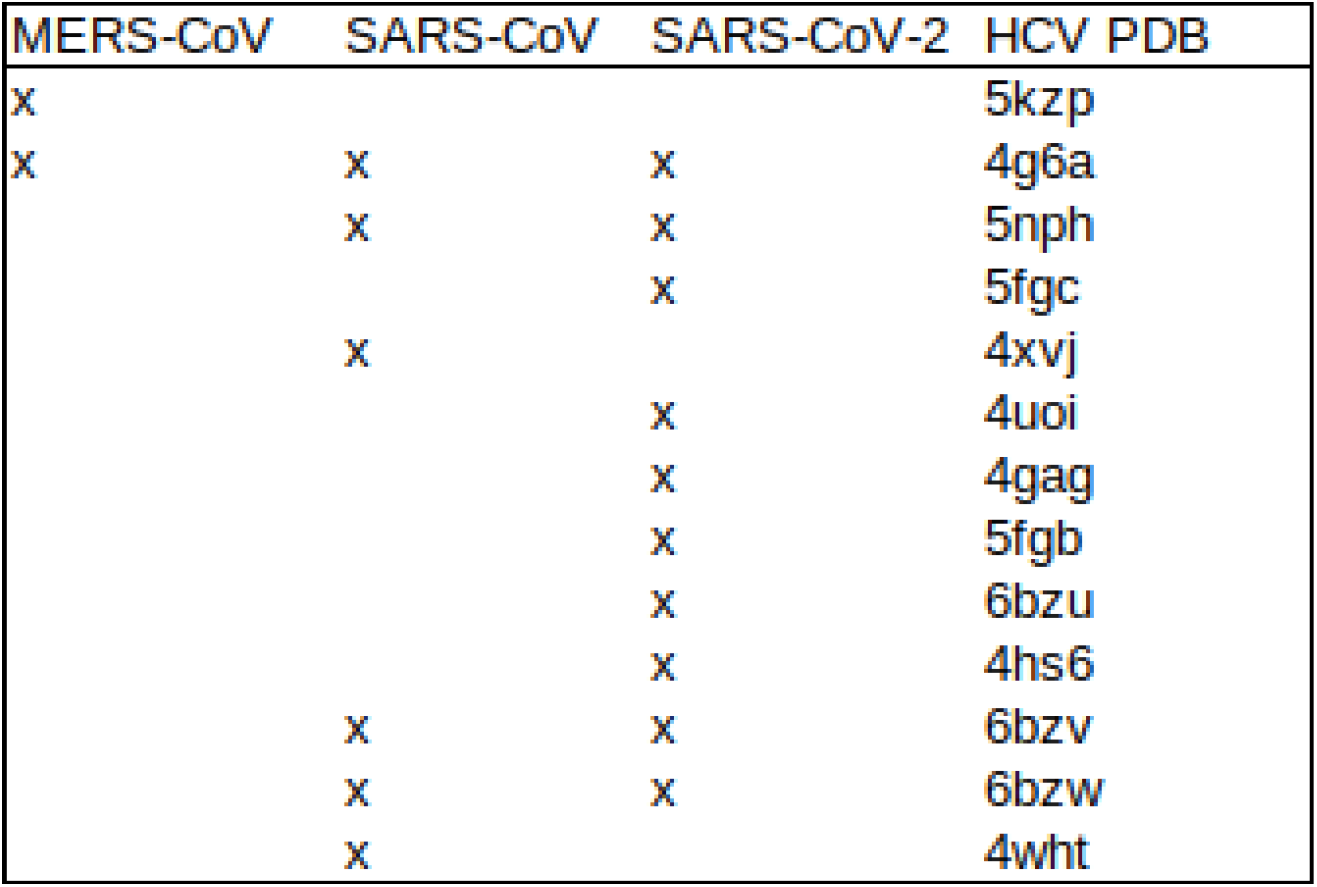
Hepatitis C virus (HCV) PDB codes returned in structural similarity screen.

## Notes

https://github.com/tlb-lab/Coronvirus-spike-mimicry

### Competing Interest Statement

The authors have declared no competing interest.

